# No cause for pause: new analyses of ramping and stepping dynamics in LIP (Rebuttal to Response to Reply to Comment on Latimer et al 2015)

**DOI:** 10.1101/160994

**Authors:** Kenneth W. Latimer, Alexander C. Huk, Jonathan W. Pillow

## Abstract

We recently presented a statistical comparison between two models of latent dynamics in macaque lateral intraparietal (LIP) area spike trains—a continuous ‘ramping’ (diffusion-to-bound) model, and a discrete ‘stepping’ model—and found that a substantial fraction of neurons (recorded in two different studies) were better supported by the stepping model (Latimer et al., 2015). Here, we respond to a recent challenge to the validity of these findings that focuses primarily on the possibility of a lower bound on LIP firing rates (Zylberberg & Shadlen, 2016). The paper in question proposed alternate formulations of the ramping model, and argued (via indirect analyses) that half the neurons in the population were better explained by the new model; if correct, this would lead to an even split in the number of neurons better explained by each model. These analyses, while interesting, do not alter the conclusions of our original paper. Here, we review the criticisms raised by Zylberberg & Shadlen and report several new analyses using models with lower bounds. First, we show that the stepping model continued to provide a better description of LIP spike trains when fit using only an early period of each trial. Second, we performed a direct model comparison between our stepping model and a ramping-with-baseline model proposed by Zylberberg & Shadlen; we found that (in a pleasing moment of agreement) roughly half the neurons were better explained by each model. Interestingly, inspection of the cells that switched classifications revealed that many did not strictly exhibit the classical ramping PSTHs that motivated these analyses in the first place. We also examined two other issues raised in recent discussions of LIP: (1) We show that a non-integrating model is consistent with some core aspects of behavioral data previously offered as evidence for continuous integration; and (2) We examine analyses based on the response covariance (“CorCE”), and show that it does not reliably distinguish ramping and stepping dynamics for our dataset. Taken together, these discussions highlight the value of data-driven characterizations of both neural and behavioral dynamics with appropriate statistical tools.

---

In Latimer et al. (2015), we compared two models of the latent dynamics governing single-trial spike trains in the lateral intraparietal (LIP) area during decision making: a continuous accumulation-to-bound or ‘ramping’ model and a discrete ‘stepping’ model. Although substantial prior literature asserted that these neurons exhibited ramping dynamics (Shadlen & Newsome, 1996; Mazurek et al., 2003; Shadlen & Newsome, 2001; Gold & Shadlen, 2007; Beck et al., 2008; Churchland et al., 2011), we performed an explicit statistical comparison of the two models and found that a substantial fraction of the neurons were better explained by the stepping model.

Recently, Zylberberg & Shadlen (2016), henceforth referred to as “ZS16” for brevity, argued that our results are the consequence of our formulation of the accumulation-to-bound model, in particular the manner in which it handles downward latent trajectories. Here we provide several lines of argument against these claims, and also take this informal opportunity to explore related topics that have arisen in discussions after publication of our original study.

Our manuscript is organized as follows. In Sec. 1, we review the original literature that supports the formulation of the accumulation-to-bound model we used in Latimer et al. (2015). In Sec. 2, we compare the stepping and ramping models on spike trains taken from an early window of the evidence-accumulation period. In Sec. 3, we discuss the simulations presented by ZS16 of a ramping model augmented with a positive baseline rate. In Sec. 4, we report a new comparison between the original stepping model and the augmented ramping model. In Sec. 5, we report a simulation analysis of a second alternative ramping model discussed by ZS16, in which which downward ramping paths are “stopped”, not at a fixed firing rate, but by an opposing ramping process. We then turn to two other issues that have arisen in recent debates, but which are not directly related to the model comparison issues considered above. In Sec. 6 we describe an exercise in which we show that non-integrating (i.e., “high threshold”) model captures many aspects of the behavioral performance in a task typically framed as the definitive example of continuous integration of probabilistic evidence (Kira et al., 2015). Lastly, in Sec. 7, we show that the stepping and ramping models fit to LIP responses appear similar when analyzed with complementary statistical approach based on correlations of the inferred spike rate (“CorCE”).

Taking these points together, we reaffirm our original conclusions that a discrete stepping model provides a viable account of the latent spike train dynamics in a substantial fraction of LIP neurons compared to the classic accumulation-to-bound model; this result holds even when the latter model is extended to incorporate lower bounds on firing rate. These findings validate the usefulness of precise, quantitative frameworks for model comparison and testing, and motivate further work to formulate richer and more flexible models of latent dynamics in LIP.

## 1 The LIP Accumulation-to-Bound (LIP-AB) hypothesis

On average, an LIP neuron’s spike rate during the deliberation phase of the motion-discrimination task ramps upwards on trials when the monkey eventually saccades to a target placed inside the cell’s response field (RF; a “T_in_ choice”), and ramps down when the monkey saccades to a choice target outside its RF (“T_out_ choice”) (Roitman & Shadlen, 2002; Gold & Shadlen, 2007; Kiani et al., 2008; Churchland et al., 2008). According to the accumulation-to-bound in LIP (“LIP-AB”) hypothesis, an individual LIP cell’s firing rate represents accumulated evidence for making a saccade into the cell’s RF, while another set of (unobserved) cells is thought to encode the evidence for a saccade to the opposing target outside the cell’s RF. The two sets of cells act as competing or racing accumulators: when one of the accumulators reaches an upper bound, it triggers the decision. In our formulation of the ramping model, the spike rate of the LIP neuron followed a rectified (to ensure non-negative spike rates) drift-diffusion process with an absorbing upper bound. However, ZS16 argued that the ramping model in our study was in this instance inappropriate, because downward ramping in our model was not bounded below (or stopped) on T_out_ trials.

The literature contains many variants and extensions of the LIP-AB model, which are tailored to match the features of different datasets, and are often described in words rather than equations. We feel these differences are best addressed through formal model specification and comparison methods, and we welcome the search for richer and more accurate models of LIP response dynamics. However, we emphasize that the accumulation-to-bound model we considered in Latimer et al. (2015) accurately reflects previous literature on ramping models of LIP responses and decision making. In fact, we selected that formulation in order to match existing claims about the LIP-AB dynamics as faithfully as possible. For example, Churchland, Kiani, Chaudhuri, Wang, Pouget, & Shadlen (2011), in describing the second-order statistics of spike counts in LIP, argued that

> “The difference in VarCE for T_in_ and T_out_ indicates that the mechanism for terminating decisions resembles a threshold or bound for the winning choice only.”

Resulaj, Kiani, Wolpert, & Shadlen (2009) included post-decision accumulation for modeling behavior, stating:

> “In this model, once the initial bound has been reached and a decision made, evidence continues to accumulate until it either reaches a new ‘change-of-mind’ bound or a time deadline terminates post-initiation processing.”

We provide an extensive list of quotes from original literature in the Appendix.

Independent of these claims about fidelity to prior literature, we agree that it is worthwhile to consider a richer suite of dynamical models. In this paper, we extend our previous analyses to consider alternate formulations of the LIP-AB model that incorporate lower bounds on spike rate. In ZS16, two different implementations of lower bounds or stopping are discussed:

1. **Rectified accumulation-to-bound plus non-zero baseline**: the spike rate is given by a linear rectified function of a continuous latent drift-diffusion process plus a non-zero baseline. We will refer to this as the *ramping-with-baseline* model.
2. **Stopping induced by competing accumulator**: the latent diffusion process is stopped on “T_out_ ” trials by an unobserved “winning” accumulator associated with the anti-preferred choice. We will refer to this as the *lower-stopping* model.

In the following, we will consider each of these possible extensions to the basic diffusion-to-bound model in turn, and present several new analyses inspired by the ideas presented in ZS16.

## 2 Early evidence-accumulation activity supports stepping

The concerns of ZS16 focused on dynamics during the later portions of the motion stimulus period, putatively after some decisions had been made. We therefore sought to remove the possible effects of stopping or lower-bounding in the ramping dynamics by focusing on the beginnings of trials. We performed our model comparison analysis again using only 300 ms segments of each trial (200-500 ms after motion onset) instead of the entire 500-1000 ms stimulus period we analyzed originally. We found that 34 out of 40 cells were better supported by the stepping model, even when the late-in-trial portions of the spike trains had been removed from the analysis (Fig. 1). This is similar to our original result which found that the stepping model better explained the responses of 31 of 40 of the same cells. These findings indicate that our results were not dependent on late, post-decision dynamics on long T_out_ trials, as claimed by ZS16.

**Figure 1:**
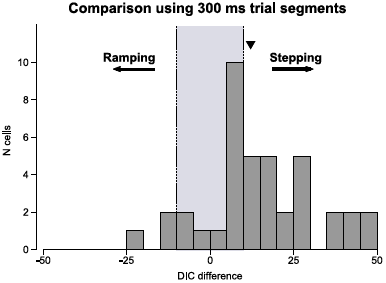
Model comparison using spike trians from 200-500 ms after motion onset for 40 LIP cells from Meister et al. (2013). The comparison is quantified as the difference in the estimated deviance information criterion (DIC) between the two models (Spiegelhalter et al., 2002) Positive difference in DIC indicate that the data were better supported by the stepping model. The black triangle denotes the median DIC difference of 12.1.

## 3 Ramping-with-baseline model

The first variant of the accumulation-to-bound model considered by ZS16 posits that neural firing rates on single trials are given by a linear-rectified function of a latent diffusion-to-bound process plus a non-zero baseline firing rate:

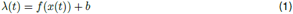

where *f* (*·*) is a rectified (or soft rectified) linear function of the latent process *x*(*t*), and *b* is an absolute lower bound, below which firing rate cannot be modulated regardless of the stimulus strength. To our knowledge, this is the first time such a rectification-plus-baseline model has been proposed for LIP.

To examine this model, ZS16 performed two simulation-based analyses. In the first analysis (Fig. 3 of ZS16), they simulated spike trains using spike rates from our original ramping model of a single idealized neuron plus a constant 10 sp/s baseline rate. They reported that the deviance information criterion (DIC) metric used in our paper favored the stepping model over the ramping model for these spike trains. We disagree with the characterization of this as a “failure to validate the diffusion model“, because these simulated data were not in fact generated by the diffusion model being tested, but were instead generated from a new model not considered in the subsequent model comparison. It is thus hard to interpret that result.

The second simulation study in ZS16 was based on a portion of the data from Roitman & Shadlen (2002), which we analyzed in our original study. They altered the data by stochastically removing spikes in order to eliminate putative baseline firing from the response rasters (using an estimated baseline firing rate for each neuron), and applied our model comparison algorithm to the resulting spike trains. This analysis is difficult to interpret for several reasons: (1) it does not explicitly define the models being tested, (2) events that actually occurred in the data (such as dips below the inferred baseline rate) can be deleted before any model comparison takes place, and (3) the stochastic data transformation adds new structure to the data (e.g., additional point-process variance). Additionally, the high baseline firing rates of some of the examples shown in Fig. 5 of ZS16 (on the order of ~50 sp/s for some cells) seem quite high for an absolute lower bound on firing, and may have exaggerated model mismatch.

Furthermore, we note that a recent study by (Kira, Yang, & Shadlen, 2015) argued against a positive lower bound on LIP firing rates in a different RT task focused on evidence accumulation in LIP:

> “The neural activity depicted in Figure 4A-that is, the neurons with the red target in RF-does not provide compelling evidence for a lower bound on T_out_ trials (i.e., green choices): the firing rates continue to reflect the accumulated evidence up to saccade initiation.”

Given the choice of dataset with a RT task, the lack of an explicit hypothesis, and a model comparison based on simulated or altered data, we feel that the analyses presented by ZS16 do not provide evidence against the results of Latimer et al. (2015). We finally note that, regardless of the concerns mentioned here, the analysis in ZS16 still classified 8 out of 16 cells as stepping, which hardly represents a reversion to the claim that LIP cells generally exhibit ramping (accumulation-to-bound) dynamics.

We also wish to make a technical comment about our fitting approach. Fig. 1 of ZS16 shows single traces denoted “hypothetical firing rate trajectories and their fits by a ramping model” to motivate the addition of a lower bound. This is a mischaracterization of our fitting method, as we did not fit firing rates on single trials; rather, we performed Bayesian inference by marginalizing (summing) over all possible latent firing rates on each trial using MCMC techniques.

## 4 Formal comparison to ramping-with-baseline model

We can include baseline firing rate as an explicit parameter in our modeling framework, and thus the lower-bound simulations in Fig. 3 of ZS16 can be formulated as a simple extension to our original ramping model. Therefore, we decided to model directly a lower bound, rather than the ad hoc approach of stochastically removing spikes from the data (and as a result, performing model comparison on simulated spike trains only). This allowed us to perform model comparison directly on real data. We added one extra parameter to the original ramping model, *b*, which gives the baseline firing rate (but did not alter the original simple stepping model). The resulting full model for trial *j* is

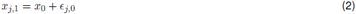

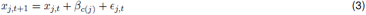

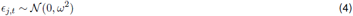

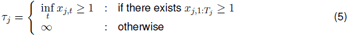

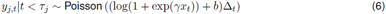

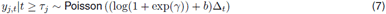

where *x* is the diffusion process, *τ*_*j*_ is the bound hit time, and *y* are the spike counts.

We analyzed the same dataset used in our original study (Meister et al., 2013) using this new rampingwith-baseline-rate model and implemented with the same Markov chain Monte Carlo methods of our original study (the posterior-mean parameters are given at the end of the appendix). We found that for these data, half of the cells were still better described by the original stepping model (Fig. 2). When we examined the PSTHs of some of the cells that switched from ‘stepping’ (as classified in our original study) to ‘ramping-with-baseline,’ we found that neither model appears appropriate for capturing the diverse range of dynamics seen in these choice-selective LIP neurons (Fig. 3). Although these cells show some form of choice-selective activity in their averaged responses, there are important differences between the average neural activity and the behavioral model of evidence accumulation, and these differences may be ignored by population-averaged statistics (see Fig. 3 caption for details). Thus, the result of a binary step-vs-ramp model comparison alone does not clearly indicate that the extended ramping model accurately describes the responses in these cells.

**Figure 2:**
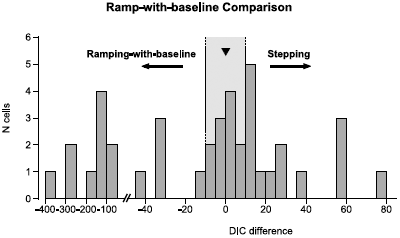
We performed Bayesian model comparison to determine whether the spike trains of LIP neurons were better explained by a ramping model with a positive baseline rate or the stepping model. Model comparison of the 40 cells analyzed in Latimer et al. with a ramping model that included a baseline firing rate. 20 of the 40 cells were still fit better by the stepping model (compared to 31/40 in the original analysis). The median ΔDIC is denoted by the black triangle. Because we kept the stepping model the same while increasing the power of the ramping model, the ΔDIC values moved only towards favoring ramping compared to the original study.

**Figure 3:**
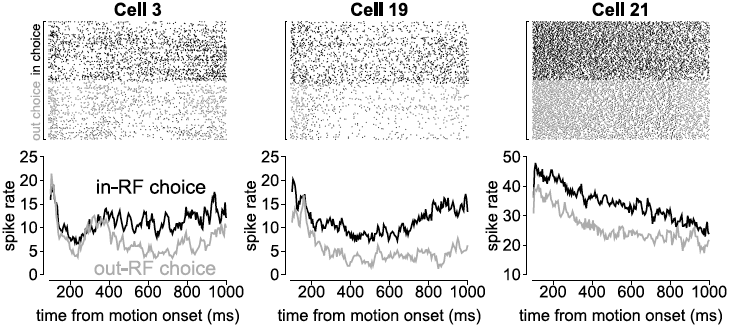
The choice-sorted PSTHs for 3 LIP cells that were classified as stepping in Latimer et al. (2015), but switched to ramping with the inclusion of a baseline rate. (left) Cell 3 exhibited nearly identical upwards ramping for T_in_ and T_out_ choices from ~200-350 ms after motion onset. The firing rates then diverge by choice, with T_in_ choices staying approximately at a constant rate. (middle) Cell 19‘s firing rate showed choice selectivity approximately 200 ms after motion onset, but the firing rate ramped down for T_in_ choices for the first ~500 ms after motion onset and then began ramping up. (right) Cell 21 showed very early choice-dependence on firing rate and the firing rates decreased throughout the decision period at similar rate for both T_in_ and T_out_ trials.

We further investigated the model fits for the cells that switched from stepping to ramping when the lower bound was included. We simulated from the ramping-with-baseline-rate model fits, and then performed model comparison on these simulations (Fig. 4A). We also performed model comparison on similar simulations from the stepping model fits (Fig. 4B). Perhaps unsurprisingly, these two models actually look quite similar (many ΔDIC values close to 0) when we consider sparse spikes only within a short window: a quick ramp up or down to a fixed level is nearly the definition of a step. In concert with this intuitive point, the difficulty in classifying this subset of the simulated data means that it can be difficult to distinguish between these versions of the ramping and stepping models within these parameter regimes for some neurons.

**Figure 4:**
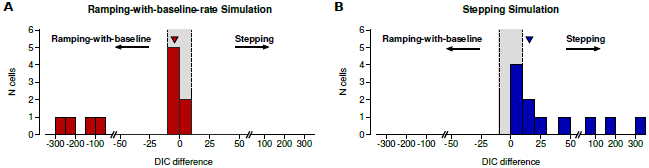
(**A**) Model comparison between the ramping-with-baseline model and the stepping model on simulations of the ramping-with-basline fits (250 trials each) to the 11 cells that were classified as “stepping” in Latimer et al. and “ramping-with-baseline” in Fig. 2. (**B**) Model comparison between the ramping-with-basline model and the stepping model on simulations of the stepping fits (250 trials each consisting of 50 trials at 5 motion coherence levels) to the same 11 cells in A. For many of these simulations, the model comparison is much closer to zero than the simulations in our original study. This suggests that the cells are operating in a regime in which continuous ramping is not meaningfully different from discrete stepping when we observe single cells. We also note that the DIC metric we chose is by no means the only way to compare models. Model evaluation is an ongoing area of research, and we encourage the exploration of a wider range of model fitness metrics for future studies (e.g., Vehtari et al., 2016).

In summary, on the timescales of the dynamics in LIP, given only sparse spikes, we do not see that a continuous, ramping process describes the data better than a discrete set of states (i.e., step-up or step-down) for many cells. For some cells, it is difficult for current techniques to distinguish between the original stepping model and the ramping-with-baseline model. We reiterate that we did not attempt to elaborate the stepping model of our original study (see Discussion for contemplation of such possibilities). These results should generally confirm the intuitions that (a) an elaborated LIP-AB results in better fits than the original LIP-AB, and (b) as stepping and ramping models become more similar, it becomes increasingly difficult to distinguish them using sparse single-trial spiking output from individual neurons. Taken together, these points do not support the notion of taking the ramping model as a strong default description of LIP dynamics.

## 5 Ramping with lower-stopping model

A second possible alternative model considered by ZS16 is one in which the firing rate on anti-preferred (“out”) choice trials exhibits “stopping” at the time that the (unobserved) competing accumulator (i.e., which integrates evidence for the other choice) hits a bound. This model has been described previously in the literature (e.g. Mazurek et al. (2003)), although the the Mazurek *et al* model actually posited stopping with a delay (relative to the time of the “winning” accumulator hitting the bound), thus allowing some post-decision accumulation. However more recent papers have relaxed or even explicitly argued against such a stopping mechanism (Ditterich, 2006a). Furthermore, evidence accumulation models with post-decision accumulation have in fact been advanced in several papers (Ditterich, 2006a,b; Resulaj et al., 2009; Fetsch et al., 2014; de Lafuente et al., 2015), including some put forth even after our study was published (Van den Berg et al., 2016a,b).

To address the effects of T_out_ stopping on our model comparison, we performed a similar exercise to the simulations in Fig. 3 of ZS16 by simulating spike rates with a stopping mechanism on T_out_ trials. We simulated two perfectly anti-correlated accumulators, each with an upper bound (Fig. 5A). The first accumulator to reach the bound would stop, and the losing accumulator would be stopped with a 100 ms delay (following the delay, *τ*_*LIP*_ given in Mazurek et al. (2003)). The spike rate was given as a rectified version of the ramping process (Fig. 5B) and spike trains were again generated from a Poisson process. To match the data from our study, the spike train lengths were sampled from a uniform distribution on 500-1000 ms. We ran our original model comparison algorithm on simulated spike trains from this new ramping process. The model comparison identified all 10 simulations as ramping (using the same model comparison code made available to ZS16), in stark opposition to the model comparison presented by ZS16 (Fig. 5C). We do not wish to oversell the impact of this simulation, because we only used one set of parameters. However, this does demonstrate that the simulations in ZS16 did not impinge upon our model comparison technique or original results.

**Figure 5:**
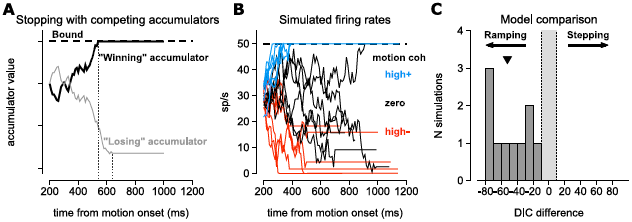
We simulated from a ramping model with a mechanism to stop downward-drifting rates. (**A**) For a single trial, we generated two perfectly anti-correlated drift diffusion rates: one represented the cell being simulated accumulating evidence for the T_in_ target and one representing accumulation towards the T_out_ target. Each process has an upper bound to signify choice, but neither is bounded below. The first accumulator to reach the upper bound (black trace) stops the diffusion process, but the losing accumulator (the accumulator that did not hit the bound; gray trace) continued to accumulate for 100 ms (Mazurek et al., 2003). (Compare to Fig. 1A of ZS16.) (**B**) Example simulated spike rates generated as a rectified samples of the process in A. Spike trains were generated as Poisson processes given the rates. The parameters used were *x*_0_ = 0.6, *ω*^2^ = 0.005, *γ* = 50, *β*_*-high*_ = − 0.02, *β*_*-low*_ = − 0.01, *β*_*zero*_ = 0, *β*_+*low*_ = 0.01, *β*_+*high*_ = 0.02. (**C**) The model comparison method from Latimer et al. consistently identified 10 simulations of 250 trials (50 trials at 5 motion coherence levels) with T_out_ stopping as ramping. The median DIC difference is denoted by the black triangle.

We agree with ZS16’s point that implementing a model comparison which includes this type of stopping for real cells poses significant challenges. This simulation was an idealized model: real single-cell responses do not always show symmetric ramping rate between IN and OUT motion, and the relative level of initial firing rate compared to the rates at the end of T_out_ and T_in_ trials show significant variability across the population (e.g., Fig. 3, and Fig. 5 of ZS16). Given the behavioral interpretation of the ramping dynamics and response heterogeneity in cortex, we do not wish to force a weighty assumption into our analyses that the dynamics underlying the decision are governed by the idiosyncrasies of the one neuron we happened to find in an experimental session. Thus, building a complete and accurate picture of single-trial dynamics requires a more complete view of LIP than population summary statistics and single-neuron recordings provide.

We also note that a model with stopping on T_out_ trials is not tightly constrained by existing single-hemisphere, single-neuron data. The hypothetical signal that is posited to stop accumulation on the observed neuron’s T_out_ trials has never been identified in these experiments, nor have the “winning” and “losing” accumulator responses been simultaneously measured and directly tied into this model, although such datasets are now possible to acquire (Bollimunta et al., 2012; Katz et al., 2016). Including such a stopping mechanism adds a great degree of flexibility to the model without evidence that the mathematical mechanism behind accumulation stopping on single trials exists in the brain.

## 6 Alternative models of the dynamics underlying behavior

The search for accumulation dynamics in LIP was motivated by the success of the drift-diffusion model (DDM) in explaining behavior in perceptual decision tasks (Shadlen & Newsome, 2001; Roitman & Shadlen, 2002; Palmer et al., 2005) Although the DDM is simple, intuitive, and has a long and rich history in cognitive psychology, choice behavior can be explained by alternate models, including models with discrete dynamics (Miller & Katz, 2010; Sadacca et al., 2016; Insabato et al., 2017). To wit, Ditterich (2006b) stated in a computational study of the data from Roitman & Shadlen that, “We will see that a relatively large class of models, both with and without temporal integration and both stationary and time-variant, turns out to be consistent with the behavioral data, including the RT distributions.” Considering such models and mechanisms can be informative. We perform one such exercise here, in no way trying to definitively prove that discrete dynamics govern decisions, but rather to point out how viable a non-integrating discrete model can be even when average behavior intuitively matches the notion of gradual accumulation.

In Latimer et al. (2015), we did not incorporate behavior into the model, preferring instead to assess models of single-trial LIP spike train dynamics that were not forced to align with decisions (a link which is especially challenging to make at the single neuron / single trial level). Given repeated references to behavior during discussions of our original results, we thought it would be constructive to consider how core attributes of the behavioral data from Kira, Yang, & Shadlen (2015) can be accounted for with various models of decision formation dynamics. This exercise suggests that alternate hypotheses should be considered for both physiology and behavior, and also highlights how the specific aspects of task design should be taken into account when considering detailed models.

This experiment used an abstract decision-making task known as the “weather prediction task”. In this task, monkeys were asked to sum up evidence given by a series of shapes (instead of motion) to pick a T_in_ or T_out_ choice. Each shape represented a probabilistic piece of evidence that the T_in_ or T_out_ choice would result in reward (Fig. 1 of Kira et al.). We considered an alternate model to evidence accumulation defined by a simple threshold (“probability summation”), in which the observer waits for a single shape with high enough evidence to trigger a decision, irrespective of previous shapes seen so far during the trial (Watson, 1979). At time each timestep *t*, the subject views a shape with evidence *x*_*t*_ and decides:

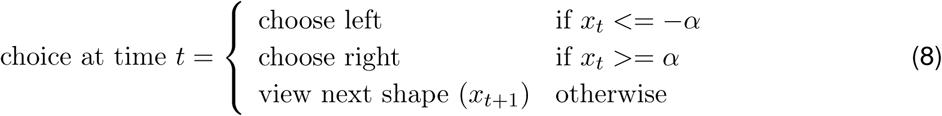

where *α* is the threshold. For simplicity, we make the additional restriction that the monkey weights each shape with the true evidence level.

Because the task included only 8 shapes with 4 levels of evidence, there are only 4 possible thresholds for the probability summation model. We found that a probability summation model with threshold parameter equal to 0.7 (i.e., the top 2 shapes for each choice) and 0.5 (top 3 shapes for each choice) accounted for both the percent correct and average number of shapes viewed for monkeys E and J, respectively (Fig. 6). This non-integrating, deterministic model captured the shape of the RT distribution for one monkey (Fig. 6B left) but not the other. This discrepancy might be explained by the fact that the monkeys were trained with different reward contingencies: Monkey J, which had a roughly exponential RT distribution (and for which the simply threshold model explained the RT distribution), received reward a fixed amount of time after making a saccadic response; Monkey E, on the other hand, was trained under more complex contingencies that motivated waiting longer before the response. As described in the original paper: “the interval to reward depended inversely on RT-a minimum of 1.8 s from onset of the first shape (e.g., immediately after validation for all RT > 1.6 s).” The paper continued to emphasize that: “Similar techniques have been used previously to counter monkeys’ natural tendency to respond quickly on choice-reaction time tasks (Hanks et al., 2014; Roitman and Shadlen, 2002).” Thus it may be difficult to distinguish simple evidence accumulation-to-bound from alternate mechanisms of evaluating (and, likely, accumulating) evidence with additional strategic components, including delaying responses based on the details of the reward contingencies (Frederick et al., 2017).

**Figure 6:**
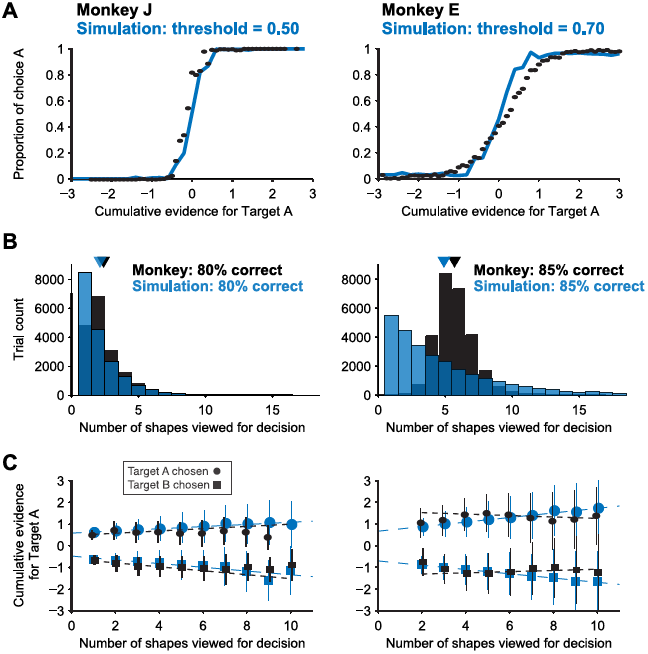
We simulated the abstract shape decision task in Kira, Yang, & Shadlen (2015) (Fig. 1A), for each trial drawing a sequence of shapes according to probability distributions extracted from Fig. 1B in Kira et al. In our simulations, the decision was made by a probability summation rule: the simulation waited until a shape with significant enough evidence was presented, ignoring all previously presented shapes. Our probability summation simulation used the true evidence-per-shape given in Fig. 1B of Kira et al. The task only has 4 levels of evidence per shape, giving only 4 possible parameter settings for the simple probability summation model. Here, we plot 2 simulations with different threshold levels (left and right, blue) overlaid with the behavioral results from 2 monkeys in Fig. 2C,E,F from Kira et al (black). The number of trials simulated was the same as the number of trials given for the corresponding monkey. (**A**) Fig. 2D from Kira et al (black) showing the percentage of target A choices as a function of the total evidence given for A compared to the probability summation model (blue). (**B**) Fig. 2C from Kira et al (black) showing the distribution of the number of shapes viewed across all trials for both monkeys are compared to the probability summation model (blue). The percent correct decisions for the simulation and the behavior are given. Because we focused here on a model with 1 parameter with only 4 possible values, we could not match these distributions exactly. However, adding additional non-integrating mechanisms in the model, like a noisy evidence threshold, can produce a closer fit (not shown). (**C**) Fig. 2E from Kira et al (black) showing total evidence at for each target at the end of the trial trial as function of the number of shapes viewed. The probability summation model (black) predicts a similar trend.

We certainly do not wish to claim that integration is not involved in this task. This exercise merely highlights the fact that an extremely simple model without integration can also accurately account for much of the behavior. Similar findings are likely in analyses of other decision making tasks, and we conclude simply that selective presentation of an integrator model (or a non-integrator model) should not by itself be taken as strong evidence of such a decision mechanism actually being at play; it should only be taken as an example of model equivalence (one of the original motivations for relying on physiology to disambiguate the decision mechanism). This exercise also highlights that the details of training contingencies should be taken into account when considering specific models of how the task is performed (Liu & Pack, 2017).

## 7 Analysis of conditional correlation (CorCE)

A separate argument put forth in support of the LIP-AB hypothesis has focused on the second order statistics of LIP spike trains. Specifically, recent work has suggested that statistics related to the variance (“VarCE”) and correlation (“CorCE”) of the conditional expectation (i.e., the firing rate) can be used to identify a signature of ramping in LIP data (Churchland et al., 2011; de Lafuente et al., 2015). Model comparison methods based on VarCE and CorCE attempt to distinguish point process noise (i.e., noise due to a Poisson-like spike generation process) from the variability of the underlying firing rate. However, many distinct doubly stochastic models (models with a stochastic latent process governing a stochastic spiking mechanism) can share similar second-order statistics (Amarasingham et al., 2015).

We examined the CorCE of simulated data generated by ramping and stepping models fit to our LIP data, and found them to be nearly indistinguishable (Fig. 7). This suggests CorCE (unlike the DIC statistic used in our original paper) does not reliably distinguish ramping from stepping dynamics because both models can produce similar CorCEs. The stepping model can produce a similar CorCE trend because the probability of having stepped up or down increases gradually over the course of the trial. This increases the spike rate variance over time just as the VarCE increases in the ramping model. Additionally, the firing rate for a single trial remains constant after the step, thereby producing an increasing CorCE along the off-diagonals through time. Therefore the stepping model is able to produce similar second order spiking statistics to the ramping model.

**Figure 7:**
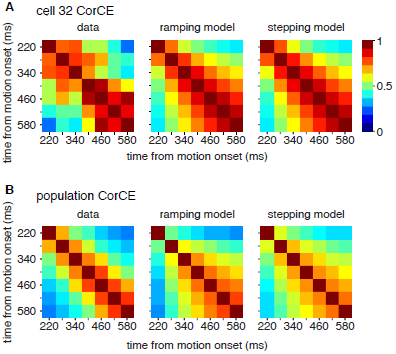
CorCE, an estimate of the temporal correlation of the spike rate relative to stimulus onset, calculated from simulated spike trains and data from Latimer et al. The CorCE was calculated in a beginning 220 ms after motion onset, and spike count statistics were computed within a centered 60 ms time bin. (**A**) CorCE for all trials of a single LIP cell (left). CorCE for simulations from ramping (middle) and stepping fits to the cell (right). The two simulations show nearly identical CorCE structure. (**B**) CorCE for the LIP population (left) and from simulations of model fits to the population for all trials. (Figure 2.28 from Latimer (2015))

The VarCE method dissects the total variance of the spike count by assuming that the point process noise is proportional to the mean firing rate, and that the remaining variance is due to the trial-to-trial variability of firing rate dynamics (in this case, those specified by the LIP-AB). However, this framework does not provide a means to assess whether this assumption holds for a particular dataset (although this can be addressed through modeling). This raises a fundamental challenge for interpreting the VarCE, because the relationship between input and Poisson spike rate is typically modeled with an accelerating nonlinearity (Chander & Chichilnisky, 2001; Miller & Troyer, 2002; Priebe et al., 2004; Jolivet et al., 2006; Goris et al., 2014). This nonlinearity implies that the contribution of point process variability may not be constant, but may instead increase with firing rate. If this phenomenon holds in LIP, it would suggest that for low levels of input (corresponding to early phases of a ramping response), the fraction of output variability contributed by the point process will be smaller; for large levels of input (corresponding to later phases of a ramping response), the point process contribution will be larger.

In the parlance of the VarCE framework, this means that *ϕ*, the point process variance contribution, may not be constant as a function of the observed changes in spike rate. Analyses that assume constant *ϕ* could thus effectively underestimate the point process contribution for larger responses (i.e., those at the end of a ramp), and erroneously assign some of that point process variance to the VarCE. This would cause the VarCE to grow with the firing rate, even if the actual input does not involve the accumulation of noise (it is merely that the statistic inherits increasing amounts of mis-assigned point process noise). The weakness of the additivity assumption has been addressed more generally by Koyama & Kobayashi (2016). Put simply, if LIP neurons exhibit forms of gain variability reported in other brain areas, VarCE may increase when the underlying firing rate increases in LIP neurons, regardless of whether it has an accumulated-noise component. We note, of course, that neither of the models we considered in our paper (ramping or stepping) incorporated firing rate variability that scaled nonlinearly with firing rate; incorporating such effects and examining their influence on model performance therefore represents one promising direction for future research.

## Conclusions

Here, we have explained the continued viability of alternates to the ramping model accounts of LIP activity, and extended them to analyses of both behavior and higher order statistics of spike trains. Specifically, we clarified that our original formulation of the LIP-AB was consistent with current literature, and also explained how the ramping-with-baseline variant of the LIP-AB presented by ZS16 was not implemented consistently with the mechanism they described, was not applied to actual LIP data, and was not evaluated within a standard model comparison framework. These issues lead us to conclude that it is difficult to interpret the original results of ZS16, and therefore to disagree with their framing of their study’s implications. In our reply, we instead implemented the ramping-with-baseline model as described by ZS16, and reported its performance in direct application to data, and in standard model comparisons of simulated data. We found that, while some cells originally classified as stepping changed to ramping with the addition of the baseline rate parameter, the resulting comparison between the original stepping model and the improved ramping model yielded an even split between cell assignments, and many of the changed-assignment cells did not even show canonical ramping dynamics in their PSTHs. Furthermore, a simpler analysis designed to mitigate contributions of post-decision lower bounds, focused only on early portions of trials, did not much affect our initial identification of a larger proportion of stepping versus ramping cells. We believe a fair summary of the state of step-ramp comparisons is that stepping remains a simple and viable model of cell dynamics, that many variants of the ramping model exist and that some of these variants can closely match the original stepping model, and that detailed analysis of single neuron responses are a reminder that neither model is a nearly-complete account of the dynamics of many individual neurons.

Additions to the LIP-AB framework, such as the T_out_ stopping put forth in ZS16, or Shadlen et al. (2016)’s suggestion of variable start times for accumulation, are certainly worth considering. Although we did not include variable start times because previous LIP studies assumed a common start time (Shadlen & Kiani, 2013; Latimer et al., 2016), we agree that it is important to consider how and when decision processes are initiated, although it is currently unclear whether behavior or physiology could (or should) be leveraged to make these single-trial constraints. Additionally, one could include collapsing bounds or urgency signals (Churchland et al., 2008), hazard rate signals (Janssen & Shadlen, 2005; Premereur et al., 2011), and changes of mind (Resulaj et al., 2009; Kiani et al., 2014; Van den Berg et al., 2016a,b). These many possibilities reflect degrees of freedom in choosing which model to pick, which challenges the ability of the resulting model to generalize across changes in the experimental paradigm or neuronal samples. Performing these sorts of single-trial statistical model comparison forces the analyst to make these particular choices in models explicit before fitting the data. Such a process also engenders detailed and precise debates of this sort, facilitated by public code sharing.

Although including a baseline rate in the ramping model improved the model fit for several cells, we could similarly improve the stepping model with additional parameters. For example, we could include trial-to-trial gain fluctuations (Goris et al., 2014), allow more than 1 transition between the 3 states (Miller & Katz, 2010; Bollimunta et al., 2012; Kiani et al., 2014), and/or include more elaborate transition dynamics. While we believe that discrete neural dynamics are potentially a part of decision making (Sadacca et al., 2016; Insabato et al., 2017), we do not wish to dive down a partisan steppingvs-ramping rabbit hole: Instead, our goal is building better models that can accurately represent the data. And although starting with a principled, binary statistical comparison seemed a noncontroversial way to begin, it may ultimately be more fruitful to use these tools to search through a larger agnostic model space. This framework could support analysis of larger-scale recordings or more flexible formulations of dynamics that could surmount the difficulties in distinguishing the stepping model from the ramping-with-baseline model for some of the individual cells.

We have focused on the dynamics of LIP spike trains, not the link of these dynamics to decisions. ZS16, however, proposed that LIP activity follows diffusion-to-bound dynamics and the continuously valued spike rate is the decision variable (accumulated evidence). We note that the role of LIP in perceptual decision making is not clearly established, as inactivating LIP during motion discrimination tasks does not clearly affect performance of the task (Katz et al., 2016). Several areas besides LIP show similar ramp-like average activity during perceptual decisions (Kim & Shadlen, 1999; Horwitz & Newsome, 2001; Ding & Gold, 2011; de Lafuente et al., 2015). Obtaining evidence that LIP is truly a direct neural correlate of DDM-like evidence accumulation is difficult precisely because the DDM captures much of the behavior, and LIP activity does appear tightly tied to the motor response that signals the decision. With decision and action tightly linked, it may not be particularly surprising that single-neuron activity in parietal cortex could show some basic parallels to the one-dimensional dynamics in the DDM, even if the neural dynamics are distinct from the DDM. More generally, we wish to put forth that a mathematical/psychological model of decision making providing a good description of the behavior does not necessarily require that the brain implements the computations described by that algorithm in spike rates of individual neurons on single trials (Heitz & Schall, 2013). Our statistical modeling results, taken together with recent findings that LIP inactivation does not change behavior, suggest that LIP activity may reflect a variety of sensory and motor factors implicated in visual-oculomotor tasks, as opposed to solely playing the role of an evidence accumulator at the level of single-cell dynamics.

Obviously progress in understanding the neurobiology of decision making will arise from a broader view than single LIP cells’ assignments in a step-vs-ramp model comparison. In one sense, we regret that the original study was taken as a rejection of the larger LIP-AB model, as even in the original paper we explicitly stated that multi-neuron recordings would be critical for pinpointing the relation between LIP’s dynamics and perceptual decision making, as the majority of models tacitly or explicitly assume population coding. It would not be surprising if individual LIP neurons exhibit more discrete dynamics than the aggregate single-trial response of a relevant population, and thus the major impact of the original study should have been to motivate efforts to understand relations between the functional properties of individual cells and the resulting circuit dynamics– as opposed to a competition between two extreme models applied to one neuron at a time.

Despite our disagreements, however, we are pleased that this debate has occurred, as it has constructively served to identify the range of models under consideration, and has confirmed the usefulness of developing and sharing tools and code for pinning these models down and performing comparisons. We are grateful that Zylberberg & Shadlen downloaded and applied our code to explore these issues. These discussions have also highlighted how a wider range of experimental paradigms will be useful for precisely controlling evidence on single trials, how training contingencies should be taken into account when building models of decision processes, and of the value in testing the generality of the associated frameworks across studies.

Our original study put forth just one pair of models and a small number of principled methods for model fitting and comparison. The continued development of more powerful statistical tools will better explore how a diverse population of cells (or brain areas) might contribute to decisions through the application of concrete, but more flexible models. The rigorous application of such statistical modeling will help determine the limits of the conclusions that can be made given the available data, which is an especially important prospect given the accelerating scope of electrophysiological recordings. We hope that the tools and approaches we have put forth, as well as the conclusion of this debate, will play a part in moving systems neuroscience beyond the search for neural correlates towards a richer biological and mechanistic understanding of cognition.

## Appendix A: Previous statements about LIP-AB model

The existence of continued integration within the DDM framework was clearly put forward in Resulaj, Kiani, Wolpert, & Shadlen (2009):

> “We proposed that the unused information could be processed after the brain has committed to an initial choice, thereby requiring an extension of the bounded-diffusion mechanism that includes post-initiation processing.”
>
> “The brain exploits information that is in the processing pipeline when the initial decision is made to subsequently either reverse or reaffirm the initial decision.”
>
> “This experiment therefore tested whether our suggested framework generalizes to a situation in which the time of the initial choice is determined by an exogenous cue. The results of this experiment, which are summarized in Supplementary Figs 1–3, confirm the finding that subjects base their initial choice on early evidence but can avail themselves of additional evidence in the processing pipeline to revise this choice. These data also conform to a variant of the bounded-accumulation mechanism with post-initiation processing”

Fetsch, Kiani, & Shadlen (2014), in a review that described a relationship between LIP and confidence in decision-making tasks, stated that

> “evidence in the processing pipeline continues to accrue favoring rightward, and if the decision is reported with a nonballistic movement (i.e., a reach of the arm), this additional evidence can cause a revision of the initial movement trajectory toward the correct choice.”

de Lafuente, Jazayeri, & Shadlen (2015) remarked in the fixed-duration setting that

> “Decision termination does not necessarily imply termination of evidence integration by neurons (Mazurek et al., 2003; Resulaj et al., 2009).”

In another study on confidence and changes of mind, Van den Berg, Anandalingam, Zylberberg, Kiani, Shadlen, & Wolpert (2016a) used a racing accumulators model with no lower bounds to conclude that

> “In the post-decision period, the accumulation therefore continues. Changes of confidence and/or decision are determined by which region the log-odds is in after processing for an additional time.” “It appears that the brain processed additional information that had already been detected but did not have time to affect the initial choice.”

Additionally, the firing rates of parietal neuron were connected to the decision variable and post-decision accumulation inVan den Berg, Zylberberg, Kiani, Shadlen, & Wolpert (2016b):

> “The accumulation is represented by neurons in the association cortex such that their firing rate is proportional to the accumulated evidence for one choice versus the other. This representation, termed a decision variable, is compared to a threshold (i.e., bound), which terminates the decision process, thereby establishing both the choice and decision time. The latter corresponds to the measured reaction time, but there are processing delays that separate these events by enough time to allow for a dissociation between the state of accumulated evidence used to terminate the decision and the evidence used to support subsequent behaviors, including a change of mind.”

Thus, continued integration in the losing accumulator was an essential facet of recent studies on confidence and change-of-mind by Shadlen, Zylberberg, and colleagues.

Finally, we note that in an earlier computational study of the data from Roitman & Shadlen (Ditterich, 2006a), the author found that

> “It turned out that the LIP data could only be reproduced when the assumption was made that the integrator hitting the boundary ceases to integrate while the other integrator continues to accumulate information.”

## Appendix B: Ramping-with-baseline parameter fits

**Table 1:**
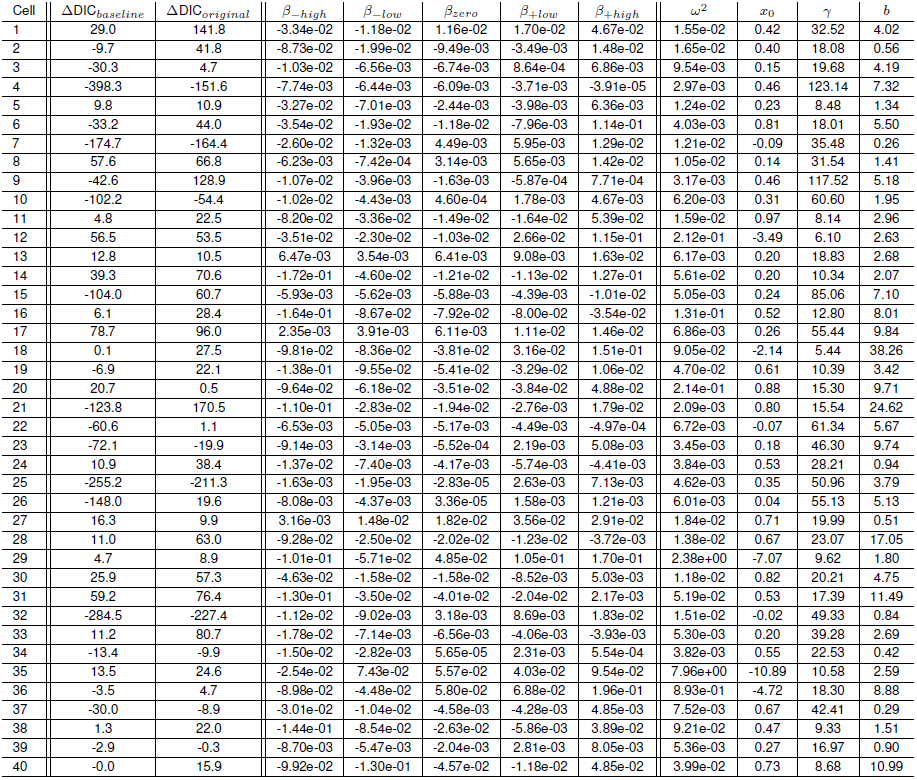
Posterior mean ramping-with-baseline parameters for the 40 cells from Latimer et al. (2015). The ΔDIC scores between the stepping model and the ramping-with-baseline model are given along with the original ΔDIC scores.

